# Human iPSC derived alveolar macrophages reveal macrophage subtype specific functions of itaconate in *M. tuberculosis* host defense

**DOI:** 10.1101/2025.07.23.664455

**Authors:** Adam Krebs, Tomi Lazarov, Anthony Reynolds, Kimberly A. Dill-McFarland, Abigail Xie, James Bean, Muxue Du, Olivier Levy, John Buglino, Aaron Zhong, Anna-Lena Neehus, Stephanie Boisson-Dupuis, Jean-Laurent Casanova, Elouise E. Kroon, Marlo Möller, Thomas R. Hawn, Ting Zhou, Lydia W.S Finley, Marc Antoine Jean Juste, Dan Fitzgerald, Frederic Geissmann, Michael S. Glickman

## Abstract

*Mycobacterium tuberculosis* (Mtb) must survive within multiple macrophage populations during infection, including alveolar macrophages (AM) and recruited inflammatory macrophages. In mice, itaconate, produced in macrophages by ACOD1 mediated decarboxylation of aconitate, has direct antimicrobial activity, modulates inflammatory cytokines, and is required for resistance to *M. tuberculosis* (Mtb) infection. The role of itaconate in human macrophages is less clear and whether itaconate mediates distinct effects in macrophage subtypes is unknown. Here, we investigated the role of itaconate in human iPSC-derived macrophages, either induced by GM-CSF to resemble alveolar macrophages (AM-Like cells), or treated with M-CSF to generate control macrophages (MCDM cells). Both types of human macrophages produce substantially less itaconate than mouse macrophages and AM-Ls produced 4-fold less itaconate than MCDMs. Surprisingly, ACOD1 deficient AM-L macrophages, but not MCDM macrophages, were permissive for Mtb growth. Moreover, itaconate functioned to dampen the Mtb induced inflammatory response in MCDMs, but not AM-L macrophages, affecting both the Type I IFN and TNF pathways. These results indicate that itaconate is involved in human macrophage responses to TB, with distinct roles in different macrophage subsets. These results also show that genetically tractable hiPSC-derived macrophages are a robust and versatile model to dissect cellular host pathogen interactions.

## Introduction

*Mycobacterium tuberculosis* (Mtb), the causative agent of tuberculosis (TB), is a human-specific pathogen that has spent millennia adapting to a cascade of immunological events to ultimately evade host control and facilitate bacterial dissemination and survival (1, 2). Mtb infection is initiated when droplets from the lung of an infected individual are inhaled and deposited in the terminal alveolus of the lung. Tissue-resident alveolar macrophages (AMs) are generally understood to be the initial host cell for Mtb, a model supported by mouse infection studies that show AMs to be the predominant infected cell at early time points after infection (3). With progressive infection, bacteria infect additional leukocyte populations recruited from the blood, including neutrophils, monocyte-derived macrophages, and interstitial macrophages (3-5). These sequential cellular hosts apply distinct pressure to *M. tuberculosis,* suggesting that the pathogen must rapidly adapt to distinct effector mechanisms in macrophage subtypes (5, 6).

Alveolar macrophages (AMs) are a distinct macrophage population that reside in lung alveoli. AMs are self-renewing tissue macrophages, largely independent from circulating monocytes, that develop during embryonic development (7-13). A hallmark feature of mouse and human AMs is the dependence on granulocyte macrophage colony-stimulating factor (GM-CSF) for the induction of the transcription factor PPAR-gamma (PPARG) which supports their development, maintenance, and function within the pulmonary environment (14-17). Both PPARG and GM-CSF deficient mice, as well as humans with genetic GM-CSF receptor deficiency or neutralizing auto-antibodies against GM-CSF, all suffer from alveolar proteinosis due to absent/defective of AMs (14, 18-22). AM have an anti-inflammatory function in pulmonary immunity via the rapid clearance of cellular debris and pathogens (13, 23-25). Macrophage dynamics in the *M. tuberculosis* infected murine lung have been described, and studies of AMs recovered from healthy human lung and infected with pulmonary pathogens like *M. tuberculosis* and influenza A virus reveal that AMs have distinct transcriptional responses compared to monocyte derived macrophages (26-29). However, little is known about the Mtb-alveolar macrophage interaction in humans. This knowledge gap is in part due to the challenge of studying AMs from humans, and the near impossibility of obtaining these lung resident cells during the clinically silent period after initial *M. tuberculosis* infection and the genetic intractability of primary human AMs.

Itaconic acid is a macrophage product implicated in macrophage-Mtb interactions. This metabolite is produced by inflammatory induction of the enzyme aconitate decarboxylase 1 (ACOD1, encoded by *Irg1*), which decarboxylates cis-aconitate to produce itaconate (30). Itaconate has complex effects on inflammation and bacterial control, including immunomodulatory effects through its inhibition of succinate dehydrogenase (SDH), prevention of IL-1β maturation, and modification of IκBζ-mediated inflammatory transcription (31-34). Itaconate can also directly inhibit bacterial growth through inhibition of bacterial isocitrate lyase and methyl-malonyl-CoA mutase (MCM) and may thereby be a direct antibacterial effector of macrophages (35). *Irg1*^-/-^ mice, which cannot produce itaconate, are highly susceptible to *M. tuberculosis* aerosol infection due to severe pulmonary inflammation (36). Murine *Irg1*^-/-^ bone marrow derived macrophages (BMDM) overexpress the TNF pathway with TB infection, but do not have a cell intrinsic defect in controlling bacterial replication, suggesting that the predominant role of itaconate in Mtb infection is the regulation of myeloid inflammatory responses rather than direct bacterial killing (36).

Murine macrophages produce very high levels of itaconate, but the role of itaconate in human macrophages is less well understood. In vitro and transfection experiments show that human ACOD1 is catalytically less active than murine ACOD1, suggesting that human macrophages may produce less itaconate (37). Although RAW264.7 cells, a commonly used murine macrophage cell line, produce ∼8mM itaconate upon stimulation with LPS, human PBMC-derived macrophages produce approximately 60µM under similar conditions (38). Beyond this difference, it is unknown whether macrophages of distinct ontogeny (i.e. tissue-resident vs blood derived) differ in production of itaconate and its role in immunomodulation and antibacterial activities.

To address these questions, we developed a genetically tractable hiPSC model of human macrophage differentiation (39-42) which can be used to produce alveolar-like macrophages (AM-L) that resemble their bona fide human AM counterparts, and syngeneic M-CSF derived macrophages as controls. We next used this model to measure and compare itaconate production in these cells, their inflammatory response and permissivity to Mtb, and the role of itaconate by generating ACOD1-deficient and isogenic control hiPSC lines. Our results show that itaconate is involved in the response of human macrophages to Mtb, despite a lower level of production in human in comparison to mice, reveal that itaconate has different functions in AM-L versus non-AM macrophages, and establish iPSC derived macrophage subtypes as a genetically tractable host cell model system to study the pathogenesis of *M. tuberculosis*.

## Results

### Generation of human alveolar-like macrophages (AM-L) through hematopoietic differentiation of peripheral blood derived induced pluripotent stem cells (iPSCs)

To develop a system for generating human alveolar-like and control macrophages, we utilized Granulocyte-macrophage colony-stimulating factor (GM-CSF) or macrophage colony-stimulating factor (M-CSF), respectively, for terminal differentiation of myeloid progenitor cells produced through hematopoietic differentiation of human induced pluripotent stem cells (iPSCs) (43). Embryoid bodies (EBs) generated from human iPSC clusters were matured for 14 days in media containing human IL-3 and M-CSF. Mature EBs produced non-adherent myeloid progenitor cells that were terminally differentiated into distinct macrophage subtypes with GM-CSF or M-CSF, which we label as AM-L (alveolar like) or MCDM(M-CSF derived macrophages), respectively (Figure S1A). AM-L macrophages survived in the absence of M-CSF (data not shown), which is consistent with the in vivo observation that, alveolar macrophages are found in M-CSF deficient mice (44-46).

Brightfield imaging and May-Grunwald Giemsa staining confirmed that both the MCDM and AM-L cells resembled macrophages based on the large, irregular shape, well-defined dark nucleus, and pale pink to purple cytosol with prominent vacuoles. (Figure S1B). To determine whether AM-L and MCDM cells differ in expression of transcription factors (TF) characteristic of AMs, we measured the mRNAs encoding PPARG, KLF4, CEBPB, and ATF5 (47). AM-L macrophages had significantly higher levels of PPARG and KLF4 compared to MCDM macrophages (Fig 1A). AM-L macrophages also have a trend towards higher relative expression of CEBPB and ATF5 compared to MCDM macrophages (Figure 1A). To further interrogate the similarity of these iPSC derived macrophage types to human alveolar and monocyte derived macrophages, we compared transcriptional profiles of iPSC macrophage to a published dataset of alveolar macrophage collected by bronchoalveolar lavage (BAL) from 6 healthy donors and monocyte derived macrophages (MDMs) from the same donors (26). Analysis of these MDM-AM pairs confirmed upregulation PPARG, KL4, and CEBPB in AMs (Figure 1B).

**Figure 1.**
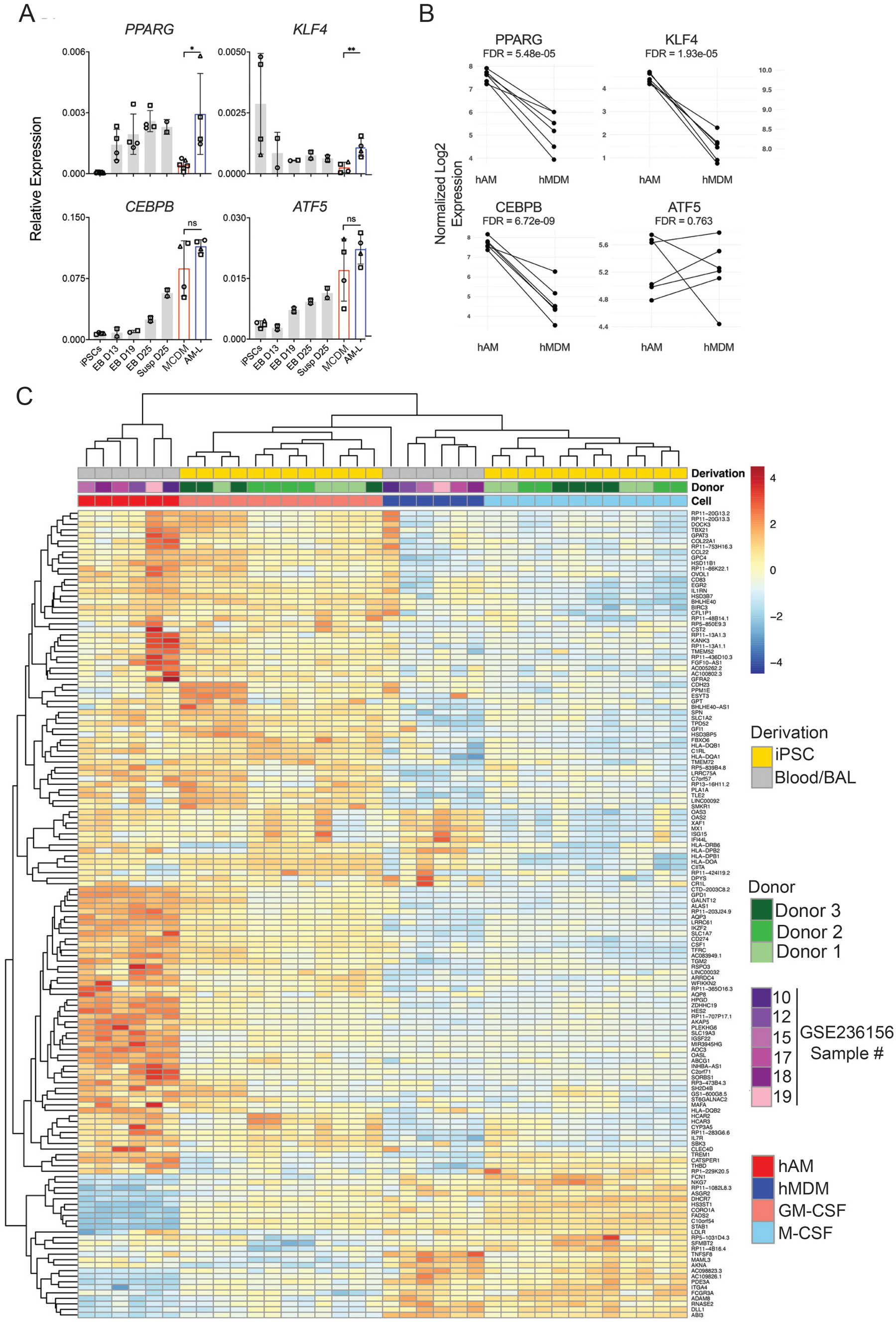
iPSC derived macrophages phenocopy the alveolar and monocyte derived macrophage transcriptional state. **(A)** qPCR assessment of transcription factors that define tissue-resident alveolar macrophages (PPAR gamma, KLF4, CEBPB, ATF5) at each stage of the iPSC differentiation process (see Figure S1A), normalized to GAPDH. Data is derived from three independent iPSC donors, each signified as a different symbol. Unpaired t-test used to assess for significant differences between MCDM and AM-L populations. ns not significant * P value < 0.05 ** P value < 0.01. **(B)** RNA levels encoding the transcription factors PPAR gamma, KLF4, CEBPB, and ATF5 in primary AMs, isolated by bronchoalveolar lavage (hAM), and monocyte derived macrophages (hMDM) derived from reanalysis of RNA sequencing data published in (26). **C)** iPSC derived AM-Ls and MCDMs resemble human alveolar and monocyte derived macrophages in basal transcriptional state. The heat map clusters the 146 most significant differentially expressed genes between iPSC derived GM-CSF (AM-L) and M-CSF (MCDM) cells from three donors (green shades) and in 6 donors (purple shades) profiled in (26). Also see Figure S2 for an expanded 736 gene set.

To more thoroughly compare iPSC derived AM-Ls and MCDMs to these primary human macrophages, we derived iPSCS and macrophages from three different human donors, performed RNA sequencing, and compared this data to ex vivo AMs vs MDMs. Using the 146 (Figure 1C) or 736 (Figure S2) most significantly differentially expressed genes between iPSC derived MCDM and AM-L cells, we clustered gene expression and observed clear clustering of primary BAL-isolated AMs with iPSC derived AMs and of MDMs with M-CSF derived iPSC macrophages (Figure 1C and S2). Taken together, these data indicate that GM-CSF dependent differentiation of EB derived human macrophages results in a gene expression profile that resembles tissue-resident alveolar macrophages and M-CSF differentiation resembles MDMs.

### Human iPSC-derived AM-L cells transcriptionally resemble of bona fide alveolar macrophages with *M. tuberculosis* infection

To assess whether human iPSC-derived macrophages resemble their ex vivo AM and MDM counterparts with *M. tuberculosis* infection, we performed RNA sequencing of AM-L and MCDM cells infected with *M. tuberculosis*. These transcriptomic profiles were compared against two transcriptomic datasets of Mtb infected macrophages (26, 39). The dataset in Campo et al. was used as a reference since it evaluates the response of primary human alveolar macrophages (AMs), isolated by bronchioalveolar lavage (BAL), to Mtb. The Arias et al. dataset was generated from Mtb infected alveolar-like macrophages differentiated from peripheral blood mononuclear cells (PBMCs) by GM-CSF, TGFβ, and synthetic surfactant (termed here blAM-L) (48) (Figure 2A).

**Figure 2.**
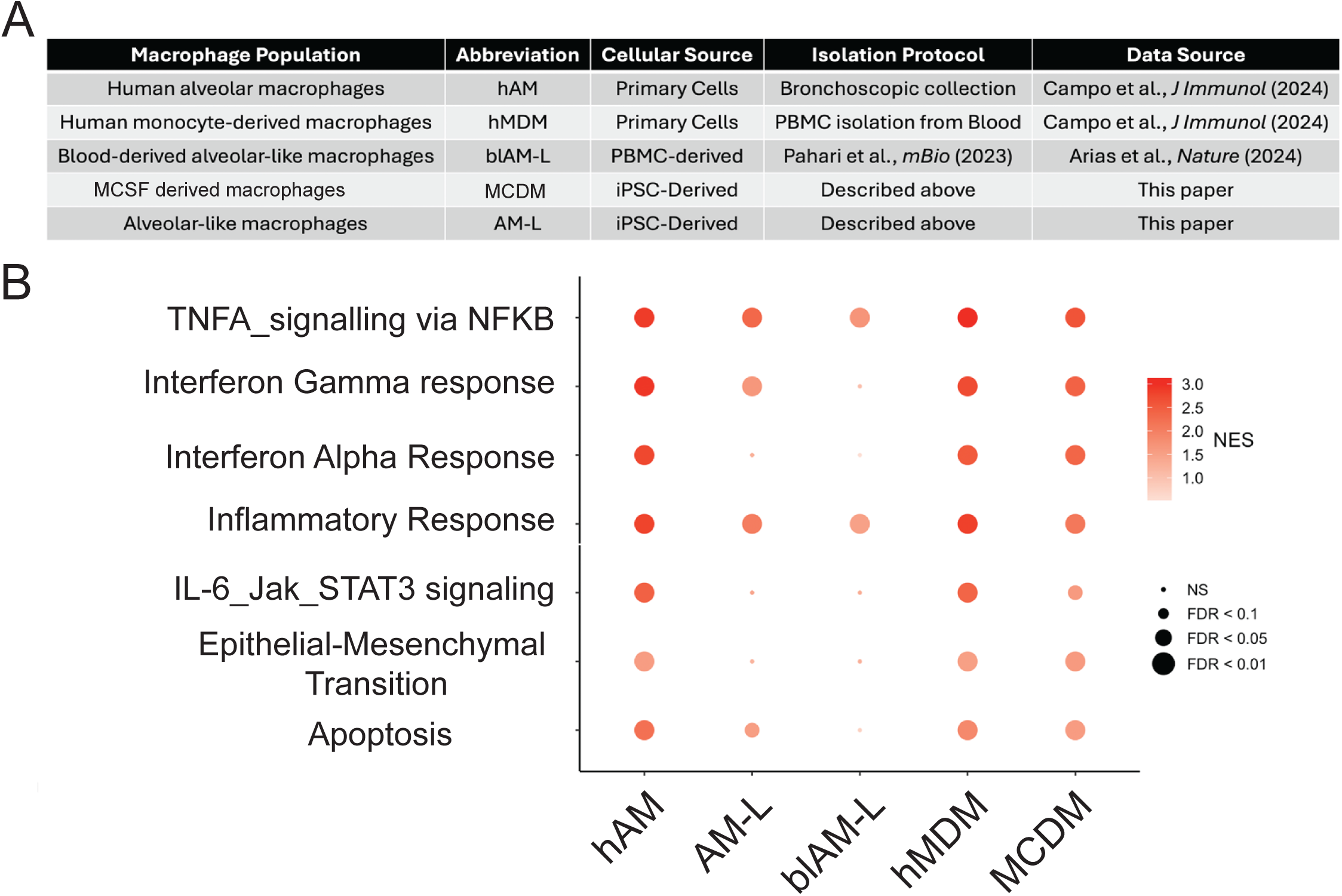
Comparison of macrophage types with *M. tuberculosis* infection. **(A)** Table of macrophage populations assessed, with abbreviations, sources, method of isolation and production, and references. **(B)** Normalized enrichment scores (NES) are plotted for each hallmark pathway induced by Mtb infection for each macrophage type listed in (A).

With Mtb infection, primary human AMs and blAM-L macrophages had a higher number of differentially expressed genes (DEGs) than iPSC derived macrophages (Figure S3). AM and blAM-L macrophages had 400 DEGs that changed in the same direction in response to Mtb, whereas 154 DEGs underwent transcriptional changes in the opposite direction (Figure S3). In contrast, all 69 DEGs shared between the AM macrophages and iPSC-derived AM-L macrophages changed in the same direction (Figure S3). blAM-L cells were the most transcriptionally reactive to TB infection and, based on the number of overlapping DEGs, more closely resembled iPSC-derived MCDM cells rather than AM-L cells.

Pathway enrichment analysis from Mtb infection induced differentially expressed genes for the three macrophage types demonstrated that all macrophages had significant enrichment in genes in the TNFα signaling by NFκB and inflammatory response pathways (Figure 2B). Genes in the interferon gamma response and apoptosis pathways were also upregulated in all macrophage types except for blAM-L macrophages (Figure 2B). Comparison of the transcriptional response of Mtb infected MDM and MCDM macrophages revealed highly similar pathway enrichment (Figure 2B). Taken together, these data indicate that the transcriptional responses of both AM-L and MCDM cells to Mtb resemble those of primary human macrophage counterparts.

### Conserved response of iPSC-derived macrophages to Mtb infection across donors

Although iPSCs, and the resulting macrophages, derived from different donors will clearly differ due to genetic heterogeneity; to serve as a model of Mtb infection, these cells should share a core response to Mtb infection independent of donor source and differentiation instance. To assess the inter-donor and differentiation variability of the iPSC macrophage model, we differentiated iPSC derived AM-L and MCDM macrophages from three different donors. PBMCs from biological siblings (donors 1 and 2) and an unrelated donor (donor 3) were reprogrammed into iPSCs and used to produce macrophages. MCDM (Figure 3A) and AM-L (Figure 3B) cells from all donors had comparable uptake of *M. tuberculosis* and MCDM cells from all donors were generally restrictive of Mtb growth such that bacterial titers did not rise substantially over the input (Figure 3A). In contrast, AM-L macrophages were generally more permissive for Mtb growth with bacterial titers rising on day 1 in all donors and further by day 3 in donors 2 and 3, with donor 3 being the most permissive (Figure 3B).

**Figure 3.**
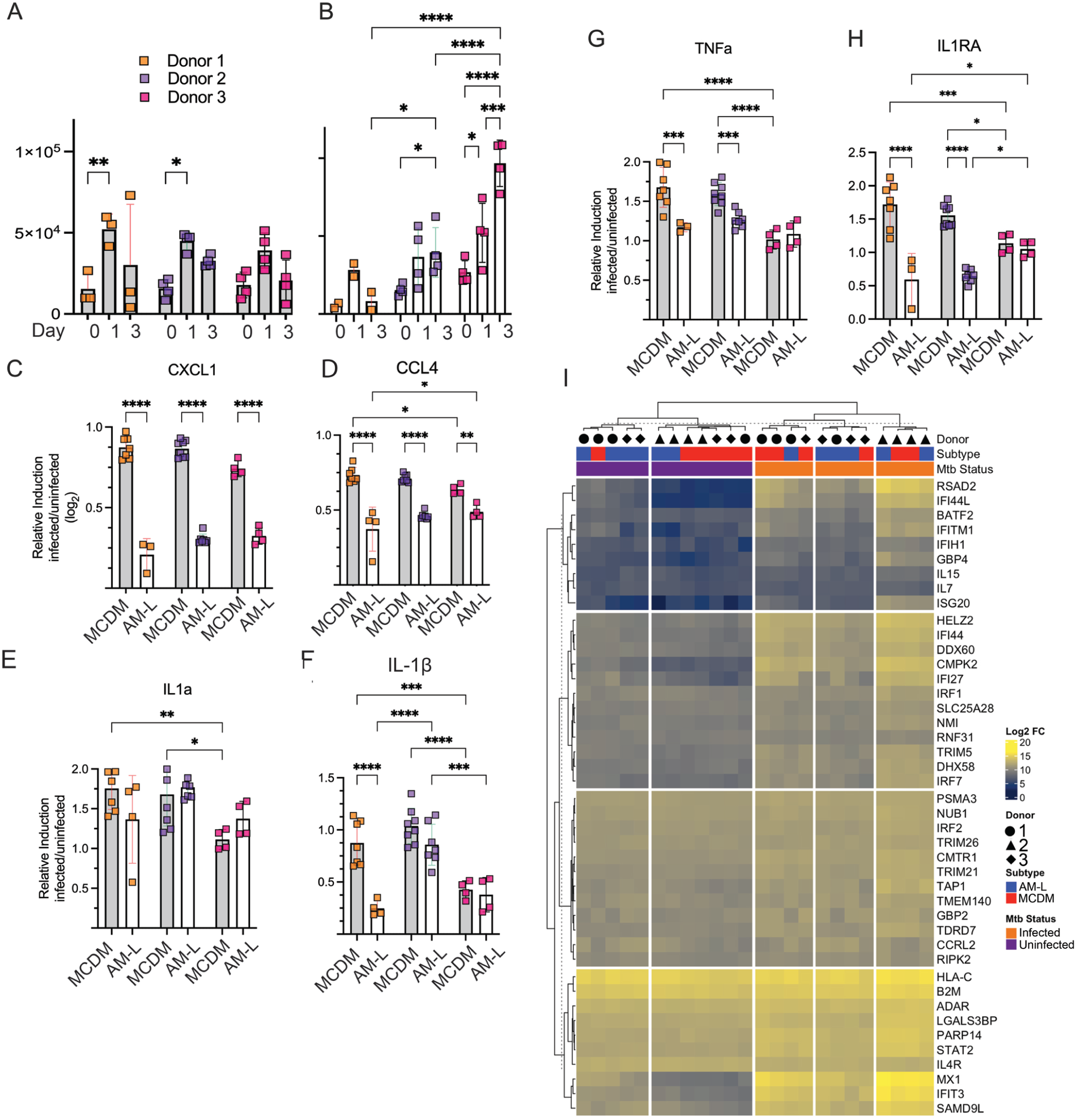
Inter donor conservation of macrophage TB responses. **(A,B)** MCDM (A) and AM-L (B) macrophages from three iPSC donors were infected with Mtb at a target multiplicity of infection (MOI) of 3. Colony forming units (CFUs) were quantified immediately post and 1-and 3-days post infection. Each donor is depicted with a different color. **(C-H) iPSC derived alveolar macrophages are hypoinflammatory** Cytokine and chemokine responses of iPSC derived macrophages on day 3 of a Mtb infection. In each graph, MCDM cells are the shaded bars and AM-L cells are clear bars, with donor identify as depicted in (A). Each Y axis is the log_2_ fold change of the indicated cytokine of infected/uninfected determined by Luminex (see methods) for CXCL-1 (C), CCL4 (D), IL-1a (E), IL-1B (F), TNF (G), and IL-1RA (H). **(I)** Heatmap depicting log_2_ fold change of hallmark type I interferon gene expression in uninfected or Mtb infected MCDM or AM-L macrophages. Partitioning is based on k-means clustering. For all panels, two-way ANOVA was used to assess variance of the samples based on infection status and within a macrophage population. * P value <0.05 ** P value <0.01 ***P value <0.0001, ****P value <0.00001.

To assess the conservation of the inflammatory response of AM-L and MCDM macrophages to Mtb infection, we measured cytokine and chemokine levels in the supernatants of infected macrophages. Mtb infection induced all measured chemokines and cytokines, regardless of macrophage subtype or donor (Figures 3C-H), with AM-L cells hypoinflammatory for most cytokines, including CXCL1 (Figure 3C), CCL4 (Figure 3D), IL-1β (Figure 3F), and TNF (Figure 3G). We observed some interdonor heterogeneity, including donor specific differences in IL-1β (Figure 3F) and IL-1RA production (Figure 3H).

To further characterize the interindividual variation in iPSC derived macrophage responses to Mtb, we performed RNA sequencing (RNAseq) of RNA from Mtb infected MCDM and AM-L from three donors. Both macrophage types from all donors robustly induced the type I interferon responsive gene set, a hallmark transcriptional response to Mtb infection, within 24-hours of Mtb infection (Figure 3I), with some weak clustering by donor identity. Overall, these data indicate that AM-L and MCDM cells consistently respond to Mtb infection largely independent of iPSC source and indicate that independent derivations from distinct iPSC sources produce macrophages with conserved responses to Mtb infection.

### AM-L and MCDM cells differ in itaconate production

Having established a human iPSC derived macrophage system to study Mtb-macrophage interactions in distinct macrophage types, we sought to use this system to understand the role of itaconate in human macrophages generally, and alveolar macrophages specifically. We first evaluated the expression of aconitate decarboxylase (ACOD1), encoded by *Irg1*, in MCDM and AM-L macrophages by stimulating with 100ng/mL of lipopolysaccharide (LPS) and immunoblotting using an anti-ACOD1 antibody. ACOD1 was not detected in unstimulated macrophages of either type (Figure 4A), but LPS stimulation strongly induced ACOD1 protein expression. MCDM macrophages produced ∼2X higher levels of ACOD1 than AM-L macrophages (Figure 4A-B). To confirm that itaconate production is dependent on *Irg1* in human iPSC derived macrophages, we generated an *Irg1* deficient iPSC line using CRISPR gene editing. Two different disruptions of exon 3 were confirmed by Sanger sequencing (Figure S4). MCDM and AM-L macrophages were generated from *Irg1^-/-^* iPSCs, indicating that itaconate is not required for macrophage development in vitro. Immunoblotting for ACOD1 in LPS stimulated *Irg1* KO macrophages of both types confirmed the absence of ACOD1 protein following LPS stimulation (Figure 4C).

**Figure 4.**
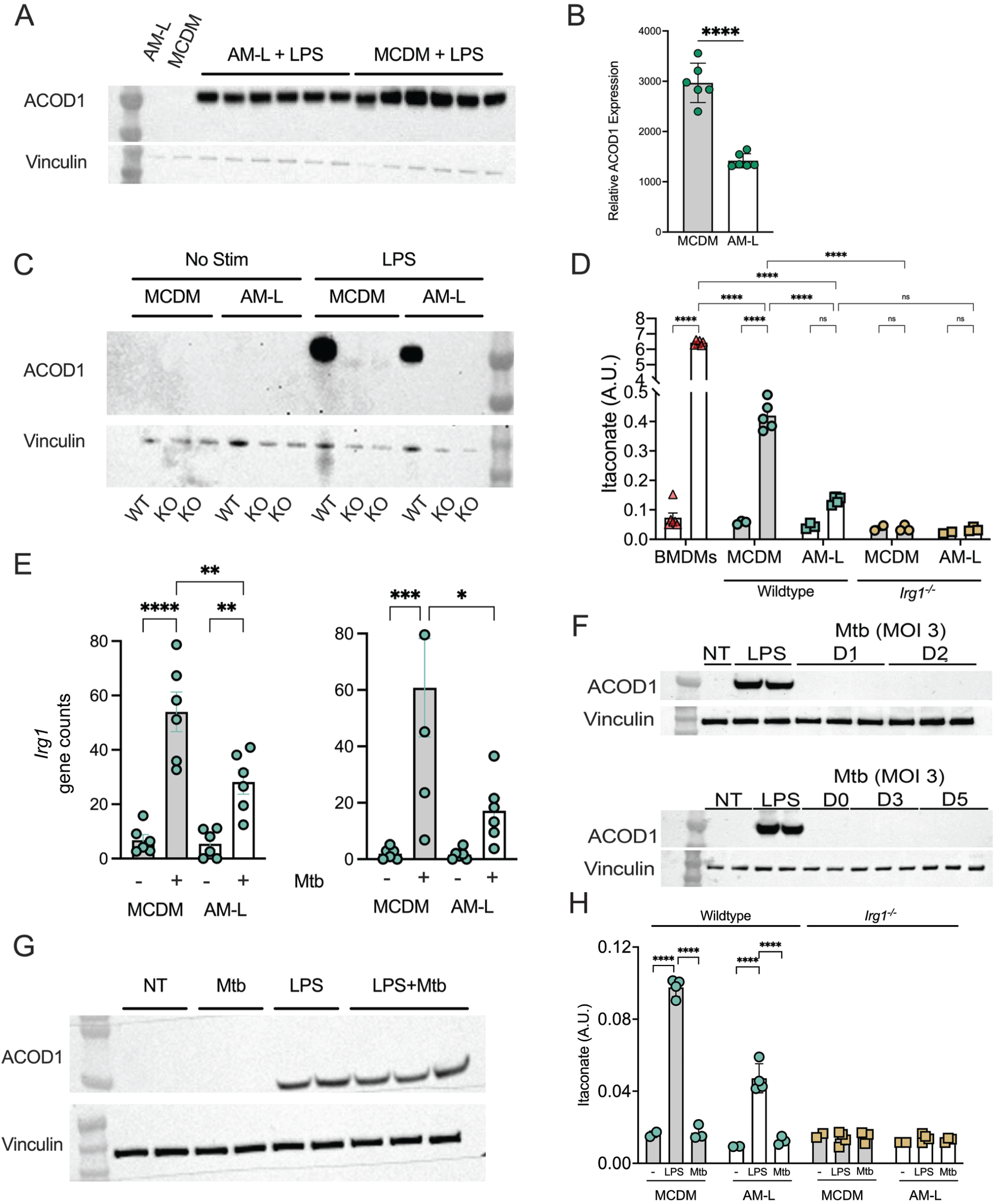
Human macrophages are feeble producers of itaconate with Mtb infection. **(A, B)** Expression of ACOD1 protein in iPSC derived macrophage subtypes. Anti-ACOD1/Vinculin immunoblot of MCDM or AM-L cells following 6-hours of stimulation with LPS (100ng/mL) (A) and normalized quantitation of ACOD1 protein levels (B). **** P value < 0.0001 by unpaired t-test. **(C) Production of *irg1*-/-macrophages**. MCDM and AM-L macrophages were differentiated from CRISPR edited iPSCs (see Figure S4) or isogenic control cells. Anti-ACOD1 immunoblot of MCDM and AM-L cells derived from *Irg1* deficient iPSCs or isogenic controls stimulated with LPS as in (A,B). **(D) Human macrophage production of itaconate**. GC-MS quantification of itaconate produced by mouse BMDMs, MCDM, AM-L, or *irg*^-/-^ MCDMs and AM-Ls following 6 hours of LPS stimulation. Itaconate levels are graphed as arbitrary units (A.U) by normalization to total protein levels. Two-way ANOVA was used to assess variance of the samples within the same treatment group and based on stimulation status. **** P value <0.0001. **(E)** Gene counts of *Irg1* mRNA following 4 hours or 24 hours of Mtb infection of MCDM or AM-L macrophages. Data is derived from three separate donors in biologic duplicates. *** P value < 0.005 **** P value <0.0001 by two-way ANOVA. **(F)** Anti-ACOD1/Vinculin immunoblot of wildtype MCDM cells following no treatment (NT), LPS stimulation (100ng/mL), or Mtb infection at MOI of 3 and assayed at days 1 and 2 (top panel), or days 0, 3, and 5 (bottom panel). **(G)** Anti-ACOD1/Vinculin immunoblot of wildtype MCDM cells following no treatment (NT), 4 hours of LPS stimulation (100ng/mL), or infection with Mtb at a multiplicity of 10 for 4 hours, or both. **(H)** GC-MS quantification of itaconate produced by MCDM and AM-L macrophages following 4 hours of infection with Mtb (MOI10) or LPS (100ng/mL). Itaconate levels were normalized to total protein levels.

To determine if the difference in ACOD1 protein found in LPS stimulated MCDM and AM-L macrophages was reflected in itaconate production, and to confirm that *Irg1*^-/-^ macrophages do not produce itaconate, we measured itaconate levels in macrophages by GC-MS. MCDM macrophages produced higher levels of itaconate compared to AM-L by a factor of 3-4-fold (Figure 4D). Itaconate levels in LPS stimulated ACOD1 deficient macrophages were similar to unstimulated cells, as assessed by GC-MS (Figure 4D). To compare human macrophage itaconate production to mouse BMDMs, we stimulated murine BMDMs with LPS and observed 20-fold more itaconate than activated MCDM macrophages and 50-fold more itaconate than activated AM-L macrophages (Figure 4D). These data demonstrate that human macrophages, despite strong induction of ACOD1 by LPS, produce significantly less itaconate that murine macrophages. iPSC derived MCDM cells produce more itaconate than alveolar like macrophages, and that CRISPR modification of iPSCs can be utilized to produce *Irg1* deficient macrophages.

### Attenuated itaconate production in Mtb infected human macrophages

LPS is a strong inducer of ACOD1 expression in both mouse (38, 49) and human macrophages (Figure 4A), but whether Mtb infection is an equivalently robust inducer of the enzyme is less clear. Prior data in murine BMDMs indicated that Mtb infection did induce mRNA encoding ACOD1, albeit to a lower level than LPS (50). We first assessed if *Irg1* transcript was induced in Mtb infected MCDM and AM-L macrophages by quantitating *Irg1* mRNA in the RNA sequencing datasets presented above and found detectable *Irg1* RNA by 4-hours of infection which was maintained for at least 24-hours (Figure 4E). Infected MCDM macrophages had a significantly higher induction of *Irg1* mRNA at both timepoints compared to AM-L macrophages, which weakly induced the transcript (Figure 4E). However, although immunoblotting with anti ACOD1 antibodies again detected expression with LPS stimulation, Mtb infection at a multiplicity of infection (MOI) of 3 did not induce detectable ACOD1 protein over either a 2 day or an extended 5-day infection (Figure 4F). Even at the non-physiologic MOI of 10, we did not observe ACOD1 protein, although LPS was still able to induce ACOD1 in Mtb infected macrophages, indicating that Mtb was not actively inhibiting expression (Figure 4G). Measurement of itaconate in Mtb infected macrophages revealed that itaconate levels did not rise above the level in uninfected cells during the first 4 hours of Mtb infection, but LPS stimulation still resulted in quantifiable levels (Figure 4H), thus paralleling the protein expression data. These data indicate that *M. tuberculosis* infection of human macrophages weakly induces the transcript encoding the ACOD1 protein and does not result in detectable itaconate production under the conditions tested.

### Itaconate deficiency impairs bacterial control by alveolar-like, but not MDM like macrophages

To assess the role of itaconate in macrophage control of *M. tuberculosis*, we assessed the capacity of *Irg1* deficient macrophages to control bacterial replication and produce cytokines and chemokines. *Irg1* deficiency had no effect on bacterial control in MCDM macrophage (Figure 5A). However, AM-L macrophages were more permissive of bacterial growth than the MCDM macrophages on day 5, and *Irg1* deficiency exacerbated this effect, indicating that this effect of itaconate on *M. tuberculosis* growth is specific to alveolar like macrophages. This result is surprising insofar as itaconate is postulated to exert a direct antimicrobial effect on intracellular bacteria, yet AM-L cells produce less ACOD1 protein and itaconate than MCDM cells.

**Figure 5.**
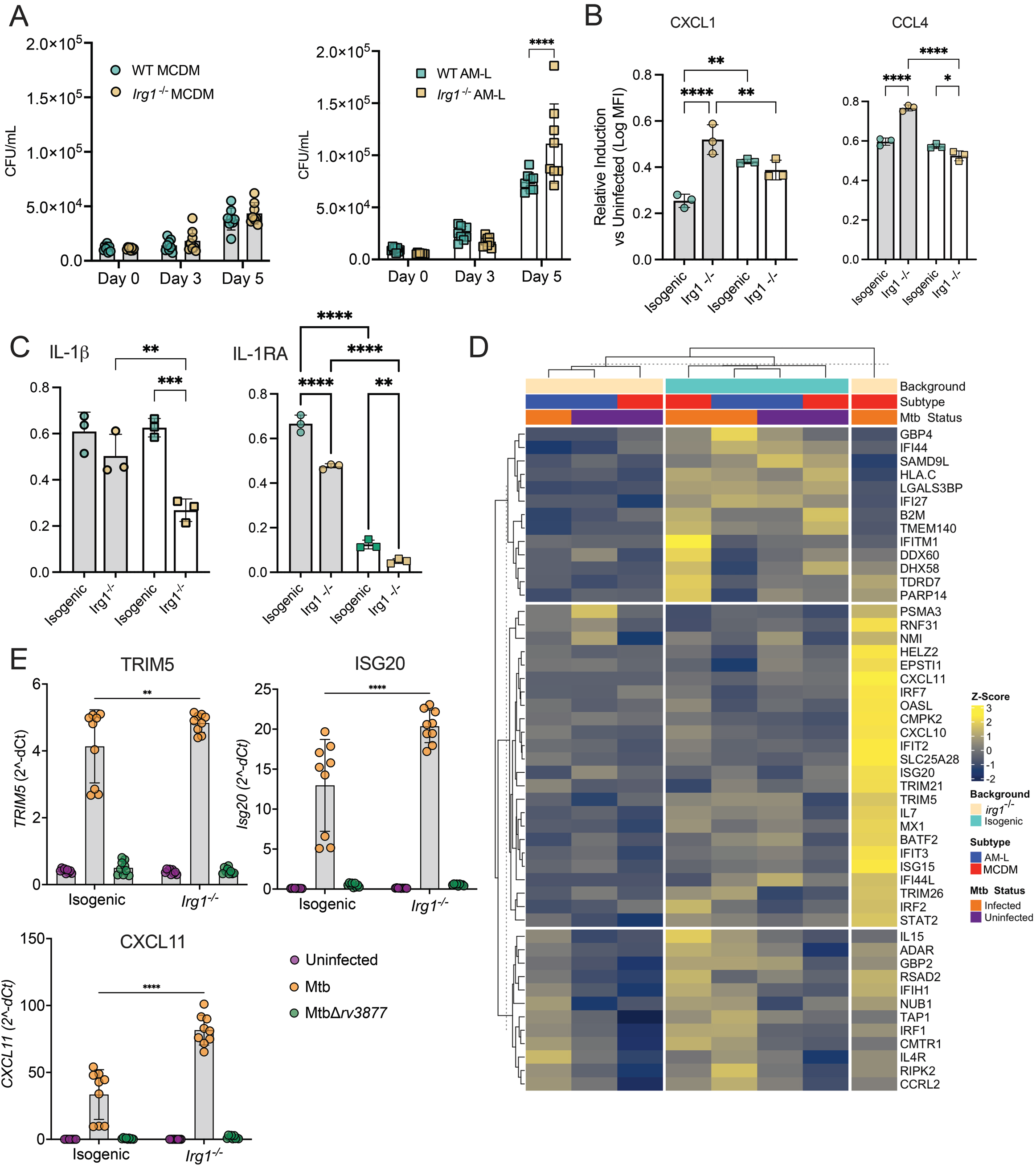
Macrophage type-specific roles of itaconate on Mtb control and inflammatory responses. **(A)** Mtb colony forming units (CFUs) quantified from wild type (green symbols) and *Irg1*-/-(gold symbols) MCDM (shaded bars) and AM-L macrophages (clear bars), infected at an MOI of 3, by day of infection **(B)** Relative concentration of CXCL1 or CCL4 produced by WT or *Irg1* ^-/-^ MCDM (shaded bars) and AM-L (clear bars) macrophages on day 3 of a Mtb infection. **(C)** Relative concentration of IL1-β and IL-1RA produced by WT and *Irg1*^-/-^ MCDM and AM-L macrophages on day 3 of a Mtb infection. **(D)** Itaconate negatively regulates the Type I interferon response to Mtb in MCDM but not AM-L macrophages. Heatmap depicting z-score of hallmark Type I interferon gene expression in MCDM and AM-L macrophages in the presence or absence of Mtb. Partitioning is based on k-means clustering. **(E)** Hyperinduction of the Type I IFN pathway in itaconate deficient macrophages is due to phagosomal permeabilization by *esx-1*. RT-qPCR quantifying *Trim5, Isg20,* and *Cxcl11* transcripts, normalized to GAPDH, in MCDM macrophages in response to infection by wildtype Mtb or MtbΔ*rv3877* (lacking the *esx-1* secretion system) for four hours. ** P value <0.01 **** P value <0.0001 by two-way ANOVA. Error bars are SD.

To verify the concentrations of itaconate required to inhibit Mtb growth in vitro, Mtb was grown in 7H9-OADC or media with propionate as a sole carbon source (7H9-ANP) supplemented with itaconate from 5µM to 15mM. 3mM itaconate in 7H9-OADC slowed Mtb growth and 15mM itaconate completely inhibited growth (Figure S5A). Itaconate was more potent in ANP media; inhibiting growth at 625µM (Figure S5A). To exclude media acidification as a cause of growth inhibition by itaconate, which has been noted by some authors (51), we buffered itaconate supplemented media with 100mM MOPS and observed that Mtb was capable of replication in 7H9-OADC even with 15mM itaconate (Figure S5B). In buffered 7H9-ANP, Mtb growth was completely inhibited by 3mM itaconate or greater and partially inhibited at 125μM, indicating that the potency of itaconate for Mtb growth inhibition is modulated by both pH and carbon source (Figure S5B).

### Macrophage subtype specific effects of itaconate on inflammatory responses to ***M. tuberculosis*.**

Since itaconate has complex effects on mouse macrophage inflammatory pathways, we determined the effect of itaconate deficiency on Mtb driven macrophage inflammatory responses by measuring chemokines and cytokines in Mtb infected macrophage supernatants. Although *Irg1* deficient macrophages produced cytokines and chemokines in response to Mtb, CXCL1 and CCL4 levels were higher in *Irg1* deficient MCDM cells than isogenic control cells, with no effect in AM-L cells (Fig 5B). In contrast, IL-1β production was lower in *Irg1* deficient AM-L cells compared to isogenic controls, with no effect in MCDM cells (Figure 5C), whereas IL-1RA required ACOD1 for full expression in both macrophage types, although MCDM cells more strongly induce this protein with infection (Figure 5C).

To better understand how *Irg1* deficiency impacts the transcriptional response of human iPSC derived MCDM and AM-L macrophages, we performed RNA sequencing after four hours of Mtb infection and compared these data to prior experiments infecting murine *Irg1* deficient BMDMs (36). We focused on the TNF and Type I IFN pathways based on the centrality of these macrophage responses to Mtb immunity (52-54), and prior reports that itaconate suppresses the TNF pathway in murine BMDMs (36). Mtb infection of wild type MCDM cells induced Type I IFN response genes (ISGs) more vigorously than AM-L cells (Figure 5D). Similarly, infection of MCDM cells broadly induced genes in the hallmark TNF/NF-Kb pathway to a greater degree than in AM-L cells, whereas AM-L cells were relatively blunted in this response (Figure S6A). In *Irg1* KO MCDM cells, transcripts in both the TNF and Type I IFN pathways were strongly overexpressed with TB infection compared to isogenic controls (Figures 5D and S6A). However, *Irg1* deficiency had little effect on either the TNF or Type I IFN pathways in AM-L cells (Figures 5D and S6A). These results demonstrate that *Irg1* deficiency in macrophages negatively regulates expression of both the TNF and Type I IFN responses with Mtb infection, but that this function is limited to MDM like macrophages and is not operative in alveolar like cells. Prior data from *Irg1* deficient mouse bone marrow derived macrophages revealed similar hyperinduction of the TNF pathway, suggesting that murine BMDMs resemble MCDM cells in response to Mtb. We reanalyzed this murine BMDM dataset, focusing specifically on Type I IFN pathway genes, and found similar overexpression of this pathway (Figure S6B), further buttressing the similarly between MCDM cells and murine BMDMs in the function of itaconate in suppressing macrophage responses to Mtb infection.

To confirm these results and determine the mechanism of Type I IFN pathway hyperinduction in *Irg1* deficient cells, we quantitated the transcripts of the ISGs TRIM5, ISG20, and CXCL11 by RT-qPCR. Mtb lacking the Esx-1 secretion system cannot permeabilize the phagosome and does not induce the Type I IFN pathway in macrophages (55-58). *Irg1*^-/-^ MCDM macrophages accumulated higher levels of TRIM5, ISG20, and CXCL11 transcripts upon Mtb infection compared to isogenic MCDM macrophages, confirming the pattern observed by RNA sequencing (Figure 5E). Infection with MtbΔ*rv3877,* a Mtb strain with a deletion of *eccD1*, a core component of the Esx1 secretion system, completely abolished the hyperexpression of these IFN target genes in both WT and *Irg1* deficient MCDM cells (Figure 5E). Taken together, these data indicate that the effects of itaconate on the macrophage ISG response proceeds through previously established pathways of phagosomal permeabilization.

## Discussion

Although *Mycobacterium tuberculosis* is an obligate human pathogen that infects via droplet deposition in the terminal alveolus of the lung, studying the initial host cell, the alveolar macrophage, is difficult due to the challenge of obtaining sufficient cells from human subjects via bronchoalveolar lavage. Even when obtained via this method, these cells are short lived and cannot be genetically manipulated. Although murine macrophage models have been widely used, less is known about how human macrophages interact with Mtb and many of these studies focus on blood derived monocyte-derived macrophages. Due to the distinct ontogeny of tissue-resident and blood derived macrophages, the antimicrobial effector mechanisms these cells apply to Mtb may be distinct. To address some of these limitations, we have developed a genetically tractable human iPSC derived macrophage model for alveolar and blood derived macrophages and used this model to examine Mtb macrophage interactions.

We found that iPSC derived macrophages differentiated with GM-CSF resemble tissue-resident alveolar macrophages based on expression of the hallmark alveolar macrophage transcription factors PPARG, KLF4, and CEBPB. Furthermore, we observed that iPSC derived AM-L cells infected with Mtb respond similarly to primary human alveolar macrophages and a newly published model of human alveolar-like macrophages derived from peripheral blood mononuclear cells (26, 39, 48). Both iPSC derived AM-L and MCDM macrophages produced chemokines and cytokines typical of Mtb infected macrophages and induced genes related to Type I interferon signaling, a hallmark of Mtb macrophage infection (52, 54). We assessed inter donor variability and generally observed conserved responses to Mtb infection across donors. These data demonstrate that iPSC derived macrophages are a useful model system for the interaction of Mtb with distinct human macrophage populations. Given the ability to derive these cells from iPSCs generated from peripheral blood, this system could also be used to interrogate clinically defined interindividual differences in TB control, especially those due to genetic etiologies manifested in macrophages (39).

Another noteworthy feature of iPSC derived macrophages is genetic tractability, which enables interrogation of specific genes to macrophage host defense, a task that is technically impractical when using primary human macrophages. We leveraged this feature to interrogate the function of itaconate in human macrophages by disrupting *Irg1*. Itaconate is an antimicrobial and immunomodulatory metabolite produced from the Krebs Cycle intermediate aconitate through the inflammatory induced expression of *Irg1*. Mouse macrophages produce mM quantities of itaconate (38), but the role of this metabolite in human macrophages is less clear. *Irg1* deficient mice, which cannot produce itaconate, are hypersusceptible to Mtb infection, but this susceptibility is due to pulmonary hyperinflammation rather than a macrophage cell intrinsic antimicrobial defect (36).

Although we found that MCDM macrophages produce significantly more itaconate than AM-L macrophages, both macrophage populations produced significantly less itaconate than mouse BMDMs, despite robust induction of the ACOD1 enzyme. Itaconate deficiency also had distinct effects on Mtb control in different macrophage types. Lack of itaconate impaired bacterial control in alveolar-like macrophages, but not MCDM macrophages. In contrast, itaconate deficient MCDM macrophages controlled Mtb like wild type, but hyperinduced the Type I interferon and TNF pathways, an inflammatory response that was not observed in itaconate deficient AM-L cells. The hyperinflammatory response in MCDM macrophages, coupled with the lack of effect on bacterial control, closely resembles observations in murine *IRG1* deficient bone marrow derived macrophages infected with Mtb (36), but differs from the function of itaconate in supporting Type I IFN pathway induction in mouse BMDMs stimulated with LPS (59). A recent report examining the role of mouse BMDMs differentiated with either M-CSF or GM-CSF in *S. aureus* skin infection found opposing roles for the metabolite in suppressing or supporting TNF production, paralleling our findings in iPSC derived human macrophage subtypes (60).

Our observed differences between itaconate deficient AMs and MCDMs are surprising because MCDMs produce more itaconate than AM-Ls, yet the latter have a more dramatic bacterial control defect, whereas the higher producers (MCDMs) have a purely inflammatory effect to itaconate deficiency. Prior data from Salmonella infection of macrophages indicates that the absolute quantity of itaconate produced is a poor correlate of macrophage intrinsic microbial control due to subcellular compartmentalization of the metabolite. Macrophages that cannot deliver itaconate to the Salmonella containing phagosomal compartment are defective for bacterial control, yet this defect does not affect cytosolic bacteria and the absolute quantity of itaconate produced is not reduced, strongly indicating that cellular trafficking of itaconate is a major determinant of its direct antimicrobial effects in macrophages (61). Although we do not probe the underlying mechanisms of our cell type specific effects of itaconate, the data may suggest that alveolar macrophages channel itaconate efficiently to the phagosome to limit bacterial growth, and thereby potentially limit the inflammatory effects of itaconate which are likely mediated by interaction with cytosolic receptors (62).

## Materials and Methods

### Antibodies and cytokines

The following antibodies were used: Alexa Fluor 488 anti-human CD206 (MRC1) Ab (ThermoFisher Scientific 564855), BV650 anti-human CD14 ( BioLegend 301836), APC/Cy7 anti-human CD45 (BioLegend 304014), PE-Cy5 anti-human CD11c (BD Biosciences 561692), PE/Cy7 anti-human CD11b (BioLegend301321), anti-IRG1 antibody (Cell Signal Technologies 77510S), HRP-conjugated human anti-vinculin (Cell Signal Technologies 18799S), IgG (H+L) Goat anti-Rabbit, HRP (Invitrogen-Fisher 656120). The following cytokines were used: recombinant Human FGF-basic (154 a.a.) (Peprotech 100-18B), recombinant Human IL3 (Peprotech 200-03-100ug), recombinant Human M-CSF( Peprotech 300-25), recombinant Human GM-CSF (Peprotech 300-03)

### Sex as a biological variable

The donor PBMCs from which iPSCs and macrophages were derived were from both male and female donors (n=1 male, n=2 female).

### Hematopoietic differentiation of human iPSCs into macrophages

Generation of MCDM and AM-L macrophages was modified from the methods described in Lachmann et al. (Stem Cell Rep 2015). Human subjects gave informed consent for blood collection at the GHESKIO center Port Au Prince Haiti under IRB protocols approved by the GHESKIO and Weil Cornell Medicine IRBs (for donors 1 and 2) or by the Health Research Ethics Committee of Stellenbosch University (N16/03/033 and N16/03/033A, for donor 3). In brief, human iPSCs, that were generated by reprogrammed peripheral blood mononuclear cells (PBMCs) with Yamanaka factors, were maintained, and expanded, on CF1 irradiated mouse feeder cells (ThermoFisher A34181) in ESC media (KO DMEM (Fisher 10829018) supplemented with 10% KO Serum Replacement (Fisher 10828028)), 1% penicillin-streptomycin (Fisher 15140122), 100µM β-mercaptoethanol (Thermo Fisher 31350010), 1% non-essential amino acids (Fisher 11140050), 2mM L-glutamine (Fisher 25-030-081) supplemented with 3-10 ng/mL basic fibroblast growth factor (bFGF). To induce embryoid body (EB) formation, iPSC colonies, that were maintained in ESC without bFGF for 5 days were disrupted by treating with type IV collagenase (ThermoFisher17104019, 250U/mL), then cultured for six days in ESC supplemented with 10µM Rock inhibitor (SigmaY0503-5MG) while on an orbital shaker (100 RPM). To facilitate germ layer formation, aggregated iPSCs were manually disrupted after six hours of shaking and Rock inhibitor was diluted four-fold with fresh ESC after three days. Mature EBs were manually transferred into APEL II differentiation media (Stem Cell Technologies 5270) supplemented with 1% penicillin-streptomycin, 5% protein free hybridoma media (Fisher 12040077), 25 ng/mL human IL-3, and 50 ng/mL human M-CSF. Following two weeks of maturation, suspension cells produced by the EBs were collected and macrophage differentiation was induced by transferring these cells into macrophage differentiation media (RPMI 1640 (Fisher MT15040CV) supplemented with 10% heat-inactivated FBS and 2mM L-glutamine) with either 100 ng/mL human M-CSF, or 100 ng/mL human GM-CSF, to produced MCDM, or AM-L, macrophages, respectively. Macrophages were considered mature after being maintained in macrophage differentiation media for five days. EBs would be maintained in culture for up to 4 months once fully mature.

### Flow cytometry of macrophages

iPSC-derived macrophages were detached by treatment with trypsin (TrypLE Express (ThermoFisher Scientific; 12605-010) for 5 minutes before centrifugation at 400xg for 5 minutes. The subsequent cellular pellet was resuspended in fluorescence-activated cell sorting (FACS) buffer (PBS +0.5% bovine serum albumin (BSA) +1 mM EDTA). After blocking Fc receptors (Miltenyi; 130-059-901) at 1:10 dilution for 10 minutes, the cells were washed with FACS buffer by centrifugation at 400xg for 5 min, and immunostained in FACS buffer with the appropriate antibodies, diluted at a range of 1:50 to 1:200, incubating for 30 minutes at 4°C. Data were acquired on either an ARIA III BD flow cytometer, or a FACS Fortessa SORP, instrument and analyzed with FlowJo. Dead cells and debris were excluded from the analysis using DAPI (1 µg/ml), side (SSC-A) and forward scatter (FSC-A) gating, and doublet exclusion using forward scatter width (FSC-W) against FSC-A. At least 5000 cells were acquired for each condition.

### Cytology and imaging of iPSC derived macrophages

iPSC-derived macrophages were prepared for flow cytometry as described above. Between 1000 and 5000 cells were then sorted into 1.5ml Eppendorf tubes, that had been precoated for 2hrs with PBS 20% FBS, containing 200µl of FBS at room temperature. The sorted cells were loaded into Cytofunnels (Fisher Scientific, BMP-CYTO-DB25) attached to Super-frost slides (Thermo Scientific, 12-550-15) before centrifugation with Cytospin 3 (Thermo Shandon) at 800rpm for 10min under medium acceleration. The slides were air-dried for at least 30 min and fixed for 10 mins in 100% methanol (Fisher Scientific, A412SK-4). Methanol fixed cells were stained in May-Gru nwald solution (Sigma-Aldrich, MG500-500mL) for 10min and washed twice with UltraPure Distilled Water (Fisher Scientific, 10977-023). The cells were then stained for 10 minutes with 10% Giemsa (Sigma-Aldrich, 48900-500mL-F) diluted in UltraPure Distilled Water. Following this stain, cells were washed twice with UltraPure Distilled Water and left to air-dry overnight. The slides were mounted with Entellan New (Millipore, 1079610100) after air-drying, and representative pictures were taken using an Axio Lab.A1 microscope (Zeiss) under a N-Achroplan 100x/01.25 objective.

### Infection of macrophages

Low passage *Mycobacterium tuberculosis* (Mtb) strain Erdman was used for infected MCDM and AM-L macrophages. Mtb was grown to log phase in 7H9 Middlebrook medium supplemented with 0.5% glycerol, 0.05% Tween-80, and 10% oleic acid-albumin-dextrose-catalase (OADC) while maintained at 37°. To prepare a single-cell mycobacterial suspension, Mtb was washed by centrifugation twice with phosphate buffered saline supplemented with 0.05% Tween-80 (PBST80), followed by centrifugation at 200 x g for 10 minutes to remove aggregated bacteria. Based on the OD600 of the final supernatant, a conversion factor of OD600 of 1.0 equates to approximately 5×108 CFU/mL was used to prepare the inoculum. Macrophages were infected at a target multiplicity of infection (MOI) of 3, unless stated otherwise. After approximately 4-hours, extracellular Mtb was removed by washing once with phosphate buffered saline (PBS). Infected macrophages were maintained in macrophage differentiation media at 37°C and 5% CO2 for up to 5 days.

### Evaluation of Mtb infected macrophages

The supernatant of Mtb infected cells was filtered twice through a 0.22µm filter before storage at -80°C. Supernatants were then used to quantify cytokine and chemokine levels by Luminex, according to the manufacturer’s instructions. To assess bacterial replication control, infected macrophages were lysed by incubating with sterile water supplemented with 0.01% Triton X-100 at room temperature for 20-40 minutes before pipetting was used to mechanically lyse cells. Cell lysates were diluted with PBST_80_ and cultured on 7H10 Middlebrook agar supplemented with 0.5% glycerol and 10% oleic acid-albumin-dextrose-catalase (OADC) for three weeks at 37°C before quantification. RNA was isolated macrophages by washing once with PBS followed by lysis and RNA preservation with TRIzol, according to the manufacturer’s instruction.

### RNA isolation and RNA sequencing

Phase separation in cells lysed in 1mL TRIzol Reagent (ThermoFisher catalog # 15596018) was induced with 200 µL chloroform. RNA was extracted from 350 µL of the aqueous phase using the miRNeasy Micro Kit (Qiagen catalog # 217084) on the QIAcube Connect (Qiagen) according to the manufacturer’s protocol. Samples were eluted in 34 µL (11749) or 15 µL (15727) RNase-free water. After RiboGreen quantification and quality control by Agilent BioAnalyzer, 29-100 ng of total RNA with RIN values of 6.5-9.8 underwent polyA selection and TruSeq library preparation according to instructions provided by Illumina (TruSeq Stranded mRNA LT Kit, catalog # RS-122-2102), with 8 cycles of PCR. Samples were barcoded and run on a NovaSeq 6000 in a PE50 run, using the NovaSeq 6000 SP Reagent Kit (100 Cycles) (Illumina). An average of 59 million paired reads was generated per sample. Ribosomal reads represented 1.1-2% of the total reads generated and the percent of mRNA bases averaged 78%.

### RNA sequencing of ACOD1 deficient macrophages

After RiboGreen quantification and quality control by Agilent BioAnalyzer, 2ng total RNA with RNA integrity numbers ranging from 4.1 to 10 underwent amplification using the SMART-Seq v4 Ultra Low Input RNA Kit (Clontech catalog # 63488), with 12 cycles of amplification. Subsequently, 7ng of amplified cDNA was used to prepare libraries with the KAPA Hyper Prep Kit (Kapa Biosystems KK8504) using 8 cycles of PCR. Samples were barcoded and run on a NovaSeq 6000 X in a PE100 run, using the NovaSeq 6000 S4 Reagent Kit (200 cycles) (Illumina). An average of 34 reads million paired reads were generated per sample. Ribosomal reads represented 0.04-0.14% of the total reads generated and the percent of mRNA bases averaged 93%. RNA sequencing data has been deposited in SRA under BioProject #: PRJNA1293194.

### RNA sequencing data analysis

Differentially expressed genes (DEGs) were determined as previously described by Campo et al (26). All three data sets were re-quality filtered, normalized, and modeled together. Briefly, sequences were quality trimmed and aligned to the human genome (GRCh38) using STAR. Alignments were quality filtered and then reads were quantified using Rsubread. Counts were filtered to protein-coding genes with a minimum 0.1 counts per million (CPM) in at least three samples and normalized via trimmed mean of means (TMM) and log_2_ CPM. DEGs were defined as FDR<0.05 for Mtb versus media in cells within a single mixed effects model with interaction term blocked by donor, ∼cell+Mtb+cell:Mtb+(1|donor), in kimma (63). For comparison of baseline gene expression of AM-L versus MCDM macrophages and ex vivo AM versus MDM macrophages, normalization and differential gene expression were performed in DESeq2 (64). Gene set enrichment analysis (GSEA) was performed using gene-level estimates (log_2_ fold changes) of Mtb vs media within cell types against the Hallmark gene sets in Broad’s Molecular Signatures Database (MSigDB)(65) using SEARchways (66). Significant gene sets were defined at FDR<0.2.

### ACOD1 immunoblotting

Mature iPSC derived macrophages were treated with LPS (100ng/mL) for either four or six hours, Mtb (multiplicity of infection:10), or Mtb (multiplicity of infection:10) and LPS (100ng/mL) for four hours. Cells were washed once with PBS then treated with trypsin and incubated at 37°C for 5 minutes. Following this incubation, trypsin was neutralized with 2x volume of cold macrophage differentiation media and cells were detached by pipetting and scraping the bottom of the well. Cells were then centrifuged at 400xg for 5 minutes before the cellular pellet was resuspended cold lysis buffer (Tris-HCl pH 7.4, NaCl, EDTA, Triton X-100, Na_3_VO_4_, phosSTOP EASYPACK, Pefabloc, and EDTA-free complete protease inhibitor cocktail) and incubated for 20 minutes on ice before then centrifuged at 16,000xg for 10 minutes at 4°C. The resulting supernatant was then transferred to a new tube and diluted 1:1 with 2x loading dye (20% glycerol, 125mM Tris HCl pH 6.8, 4% SDS, 0.2% bromophenol blue, 1xDTT) and incubated at 85°C for one hour. Protein was then separated on a 4–12% NuPAGE bis-tris poly acrylamide gel. Equal loading was confirmed with commercially available HRP-conjugated human anti-vinculin antibody at a 1:1000 dilution. ACOD1 was detected with human anti-IRG1 antibody at a 1:1000. Blots were blocked and probed in 5% Omniblot non-fat milk in 1×PBST_20_ (PBS + 0.01% Tween 20). Blots were imaged with iBright FL1000 and quantification of band intensity was conducted with FIJI.

### Generation and validation of ACOD1 knockout iPSC line

CRISPR sgRNA target was designed using the web resource at https://www.benchling.com/crispr. The sgRNA target sequence: ACAAAAGCAGCATATGTGGG was cloned into the pSpCas9(BB)-2A-Puro (PX459) V2.0 vector (Addgene plasmid #62988) to make the gene targeting construct. To knockout ACOD1, 170A iPSC line were dissociated using Accutase (Innovative Cell Technologies), and electroporated using Amaxa 4D-Nucleofector (Lonza) following manufacturer’s instructions. The cells were then seeded, and four days later, iPSCs were dissociated into single cells with Accutase and subcloned. Around 10 days later, individual colonies derived from single cells were picked, mechanically disaggregated and replated into two individual wells of 96-well plates. A portion of the cells was analyzed by PCR and sanger sequencing using amplification primers (Forward-AGGGCTTCTATCTGTGGCAA, Reverse-ACTTGAGAGAAACTGGCCCC) spanning to Crispr editing site and sequenced by Sanger sequencing using primer CCTTCCATCTTCTTTCCTCTGC. Biallelic frameshift mutants were chose as knockout clones. Wildtype clonal lines from the same targeting experiments were included as controls.

### Detection of itaconate by GC-MS

Mature iPSC derived macrophages were seeded in a six-well, tissue culture treated, plate. These cells were then treated with either LPS (100ng/mL), Mtb (multiplicity of 10), or LPS (100ng/mL) and Mtb (multiplicity of 10) together for four hours. At collection, metabolites were extracted with 450µl ice-cold 80% methanol containing 2 μM deuterated 2-hydroxyglutarate (d-2-hydroxyglutaric-2,3,3,4,4-d5 acid (d5-2HG)), while maintained on dry ice. After at least an overnight incubation at −80°C, lysates were collected and centrifuged at 21,000g for 20 min at 4°C to remove protein. Samples that were treated with Mtb were then filtered twice through a 0.22µm filter. All extracts were further processed using gas chromatography coupled with mass spectrometry (GC–MS) as described below.

Metabolite extracts were dried in an evaporator (Genevac EZ-2 Elite) and resuspended by incubating with shaking at 30°C for 2 hours in 50 µL of 40 mg mL-1 methoxyamine hydrochloride in pyridine. The metabolites were further derivatized by adding either 25 µL of N-methyl-N-(trimethylsilyl) trifluoroacetamide (Thermo Fisher Scientific) and 80 µL ethyl acetate (Sigma-Aldrich), or 80µL MSTFA, and then incubated at 37°C for 30 min. The samples were analyzed using the Agilent 7890A gas chromatograph coupled to an Agilent 5975C mass selective detector. The gas chromatograph was operated in splitless injection mode with constant helium gas flow at 1mL min-1; 1uL of derivatized metabolites was injected onto an HP-5ms column and the gas chromatograph oven temperature increased from 60°C to 290°C over 25 min. Peaks representing compounds of interest were extracted and integrated using MassHunter v.B.08 (Agilent Technologies) and then normalized to both the internal standard (d5-2HG) peak area and the protein content of duplicate samples as determined using the BCA assay (Thermo Fisher Scientific). Steady-state metabolite pool levels were derived by quantifying the following ions: d5-2HG, 354 m/z; and itaconate, 259 m/z. Peaks were manually inspected and verified relative to known spectra for each metabolite.

### Determination of Mtb sensitivity to itaconate

Mtb*-*Erdman was grown in liquid 7H9 media was supplemented with 0.05% tween_80_ and either 10% OADC (oleic acid-albumin-dextrose-catalase) or, 10% ANP (albumin-NaCl-10mM sodium propionate). Itaconic acid (Sigma-I29204) was then added to these liquid medias at a range of 30mM down to 2µM. These medias were also prepared with 100mM MOPS supplementation to prevent a pH change due to the itaconate addition. Once Mtb*-*Erdman reached an optical density at 600nm (OD_600_) of 0.4-0.7, the bacterial culture was then diluted to an OD_600_ of 0.06 and mixed at a 1:1 ratio with media containing itaconate. This resulted in a final OD_600_ of 0.03 and an itaconate range of 15mM to 1µM. The OD_600_ of the culture was then assessed every 2-3 days to quantify bacterial growth.

### RT-qPCR assessment of type I interferon genes

AM-L and MCDM macrophages derived from either wildtype or *Irg1* deficient iPSCs were infected with either Mtb-Erdman or MtbΔRv3877 at a MOI of 3 as stated previously. Following a 4-hour incubation with Mtb, the media was removed, and the well was washed with phosphate buffered saline before being lysed with TRIzol. RNA was purified from the TRIzol preserved samples with a Direct-zol RNA MiniPrep (Fisher, # 50-444-622), according to the manufacturer’s instructions. cDNA was generated from these samples with a Maxima First Strand cDNA Synthesis Kit (Fisher, K1672). To quantify TRIM5, ISG20, CXCL11 induction, a TaqMan fast advance master mix for qPCR was used (Fisher, # 44-445-57). For this assay, we used predesigned primers and probes from Fisher. This assay used TRIM5 (ThermoFisher, Hs01552558_m1) conjugated to FAM, ISG20 (ThermoFisher, Hs00158122_m1) conjugated to FAM, or CXCL11 (ThermoFisher, Hs00171138_m1) conjugated to FAM. As an internal reference, GAPDH (ThermoFisher, # Hs02786624_g1) conjugated to VIC, was used. To assess for induction of these transcripts, ΔΔCT was normalized to GAPDH for each gene.

## Acknowledgements

The authors thank the donors of PBMCs and all members of the Glickman and Geissmann labs for discussions. This work was supported by U19AI135990 (Human Pathogen Mapping Initiative, U19AI162568 (Tri-I TBRU) and R01AI124349. This research was funded in part through the NIH/NCI Cancer Center Support Grant P30 CA008748.

## Supplementary Figure Legends

**Supplementary Figure 1**

**(A)** Schematic of hematopoietic differentiation of iPSCs into either GM-CSF-dependent (AM-L), or M-CSF-dependent (MCDM), macrophages. Representative bright field images of cells at each stage are shown underneath the timeline.

**(B)** Brightfield images and May-Grunwald Giemsa staining of AM-L and MCDM cell populations.

**Supplementary Figure 2**

Heat map clustering the 736 most significant differentially expressed genes between iPSC derived GM-CSF (AM-L) and M-CSF (MCDM) cells from three donors (green shades) and in 6 donors profiled in (26). Related to Figure 1.

**Supplementary Figure 3**

Upset Plot comparing differentially expressed genes (DEGs) of three different Mtb infected macrophage types. The bar graph at the bottom left corner depicts the total number of genes found to be differentially expressed (up or down) in a given macrophage type (Mtb infected/uninfected). The intersection of DEGs between each macrophage population is represented by the bar graph on the top of the figure with the groups being compared being noted by the dots connected by the lines below the x-axis.

**Supplementary Figure 4** Sanger Sequencing confirming CRISPR disruption of *Irg1* exon 3 in iPSCs

**(A)** Wildtype sequence of *Irg1* identified in isogenic controls.

**(B)** Clone 1 of *Irg1* KO confirmed to have two different deletions in allele 1 and 2, with location of deletion signified by a star on the chromatogram.

**(C)** Clone 2 of *Irg1* KO confirmed to have a single T insertion in exon 3 on both alleles, as signified by the four-pronged asterisk.

**Supplementary Figure 5** *M. tuberculosis* sensitivity to itaconate is carbon source and pH dependent

**(A)** Assessment of bacterial replication of Mtb cultured in either 7H9-OADC or 7H9-ANP with increasing concentrations of itaconic acid, ranging from 5µM up to 15mM. Quantification of optical density at 600nm was assessed over the course of 13 days. Each point represents the mean of biological triplicates.

**(B)** Bacterial replication of Mtb cultured in either MOPS buffered 7H9-OADC or 7H9-ANP with the indicated concentrations of itaconic acid from 5µM to 15mM. Each point represents the mean of biological triplicates.

**Supplementary Figure 6**

**(A)** Expression levels of TNF pathway transcripts in AM-L and MCDM WT and *Irg1* KO macrophages with and without Mtb infection.

**(B)** Reanalysis of data from Nair et al showing the type I interferon pathway in murine WT or *irg1* KO BMDMs macrophages with or without Mtb infection.

