## Supplementary material for "Human iPSC derived alveolar macrophages reveal macrophage subtype specific functions of itaconate in *M. tuberculosis* host defense": Figures S1-S6

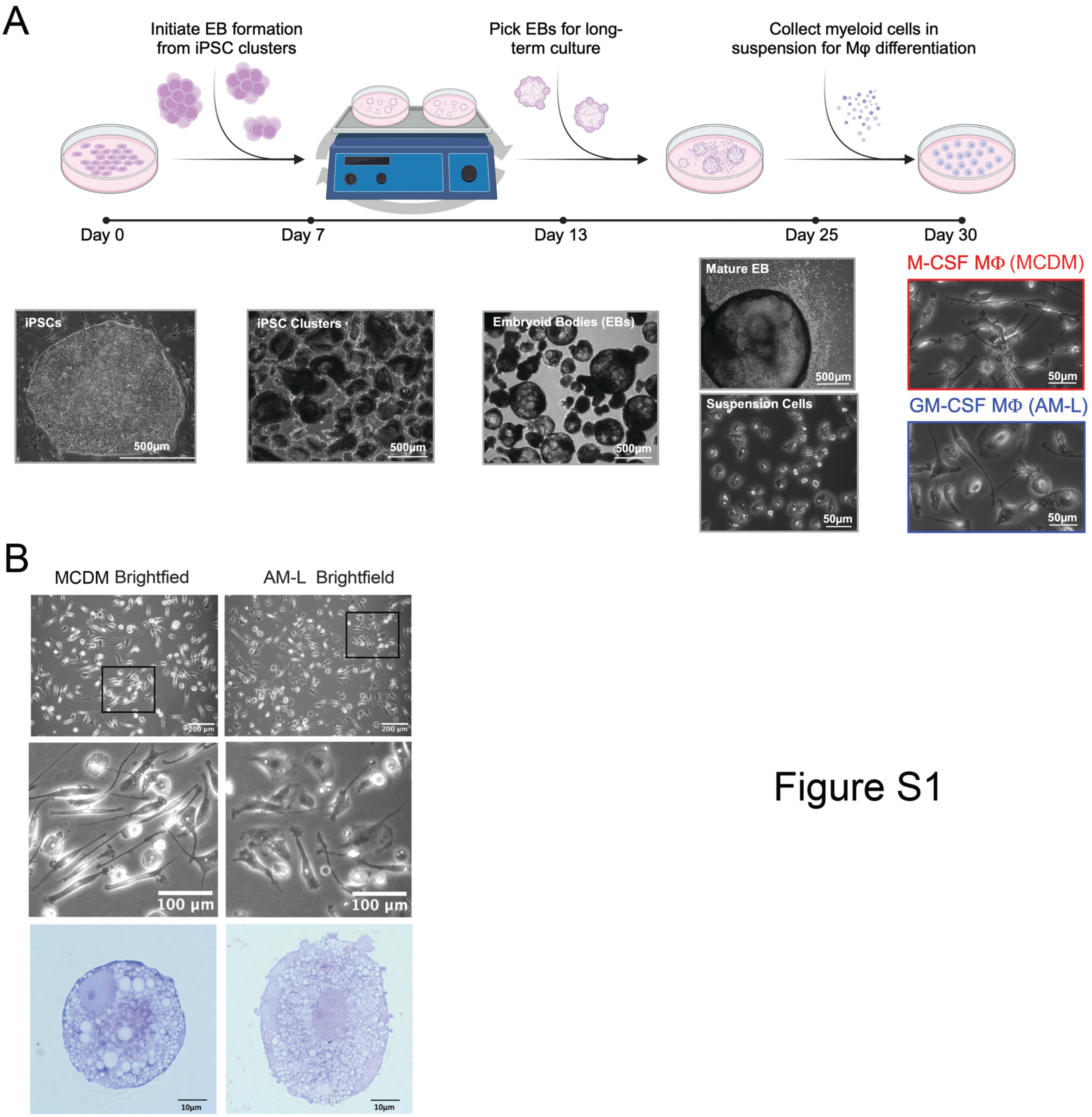
A
Initiate EB formation from iPSC clusters
Pick EBs for long-term culture
Collect myeloid cells in suspension for Mφ differentiation
Day 0
Day 7
Day 13
Day 25
Day 30
iPSCs
500μm
iPSC Clusters
500μm
Embryoid Bodies (EBs)
500μm
Mature EB
500μm
Suspension Cells
50μm
M-CSF MΦ (MCDM)
50μm
GM-CSF MΦ (AM-L)
50μm
B
MCDM Brightfied
AM-L Brightfield
200 μm
200 μm
100 μm
100 μm
10μm
10μm

Figure S1

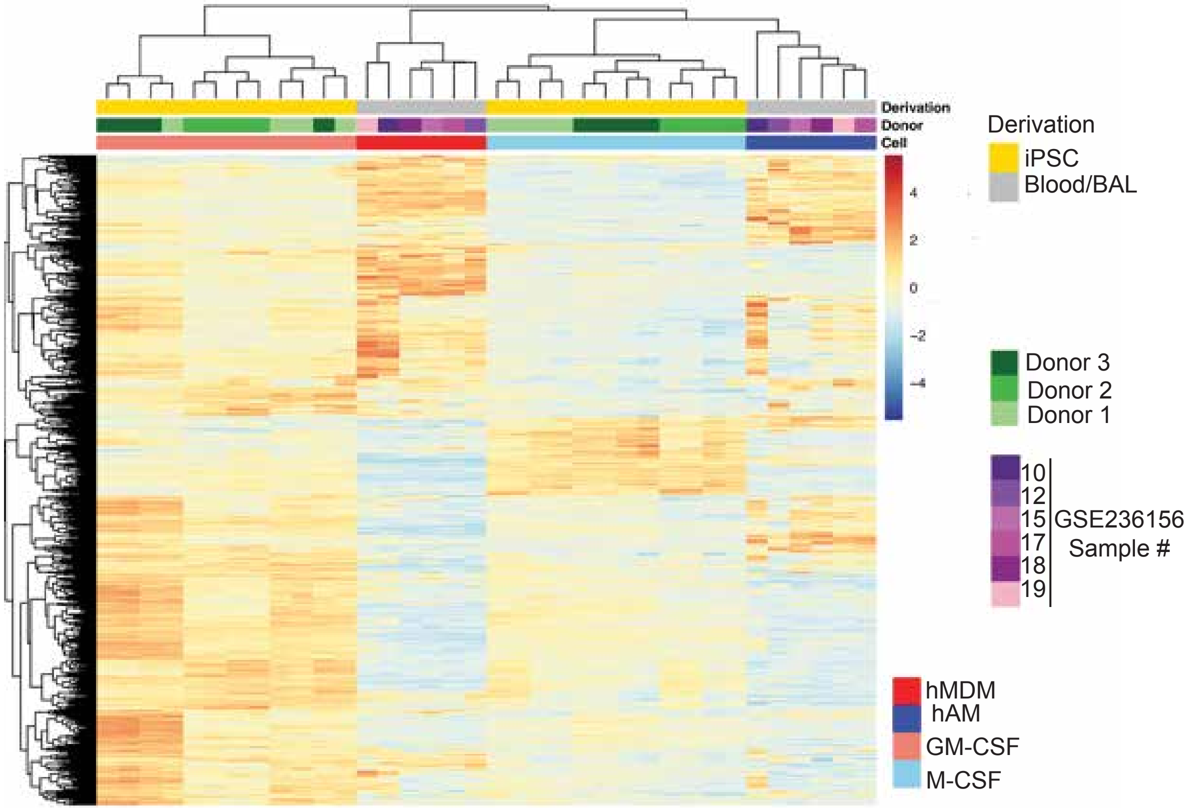
Derivation
Donor
Cell
4
2
0
-2
-4
Derivation
iPSC
Blood/BAL
Donor 3
Donor 2
Donor 1
10
12
15
17
18
19
GSE236156
Sample #
hMDM
hAM
GM-CSF
M-CSF

Figure S2

Figure S3

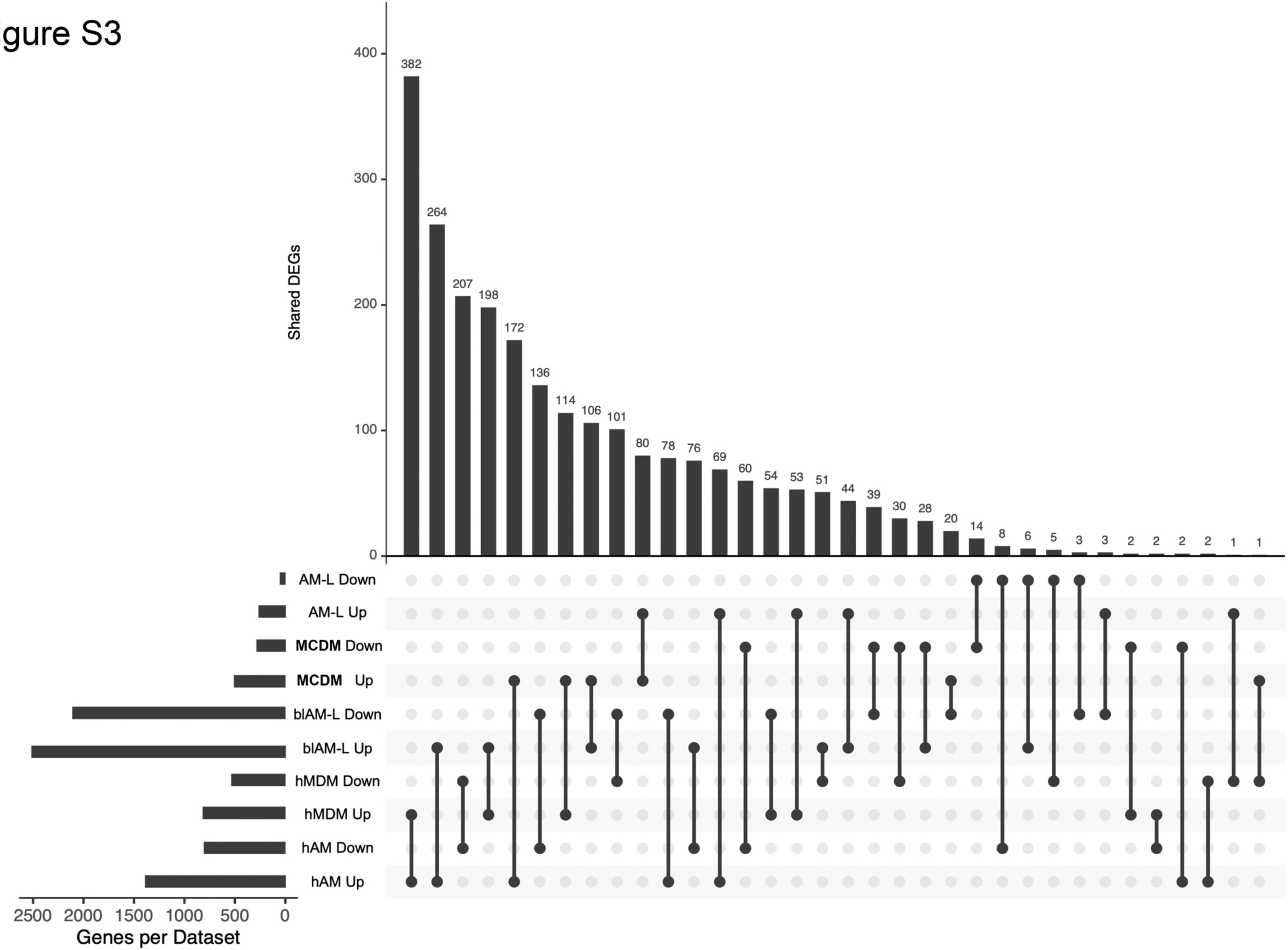
Shared DEGs
400
300
200
100
0
382
264
207
198
172
136
114
106
101
80
78
76
69
60
54
53
51
44
39
30
28
20
14
8
6
5
3
3
2
2
2
2
1
1
AM-L Down
AM-L Up
MCDM Down
MCDM Up
blAM-L Down
blAM-L Up
hMDM Down
hMDM Up
hAM Down
hAM Up
2500
2000
1500
1000
500
0
Genes per Dataset

A

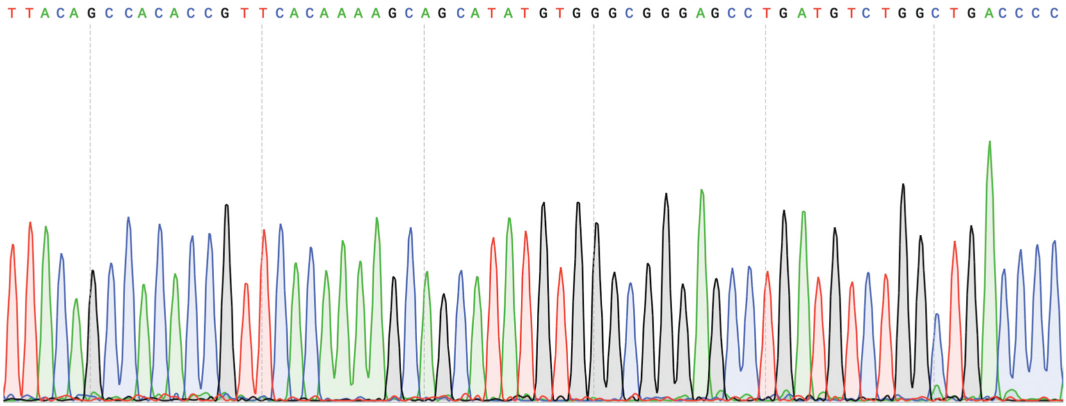


-allel 1 & 2: ACAAAAGCAGCATATGTGGGCGGGAGCCTGATGTCTGGCTGACCCC

B

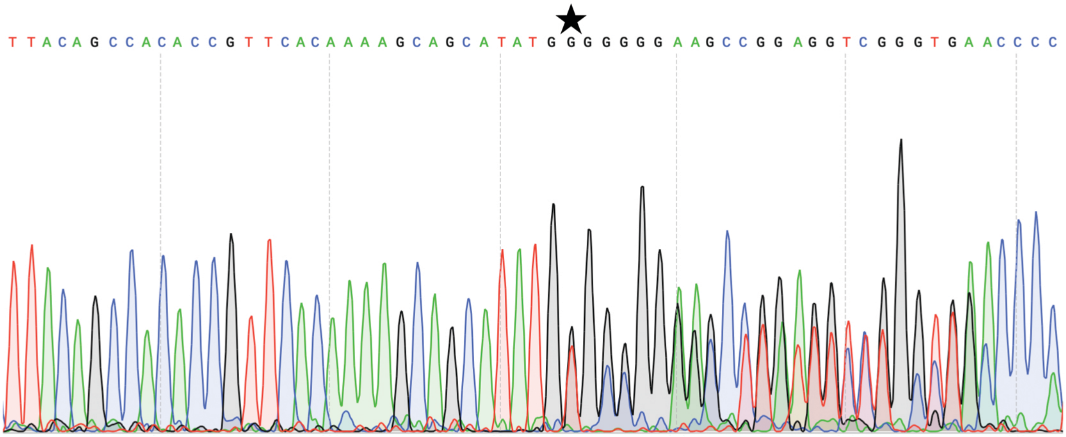


-allel1: ACAAAAGCAGCATATGTG--CGGGAGCCTGATGTCTGGCTGACCCC (Delete 2)
-allel2: ACAAAAGCAGCATATG-GGGCGGGAGCCTGATGTCTGGCTGACCC (Delete 1)

C

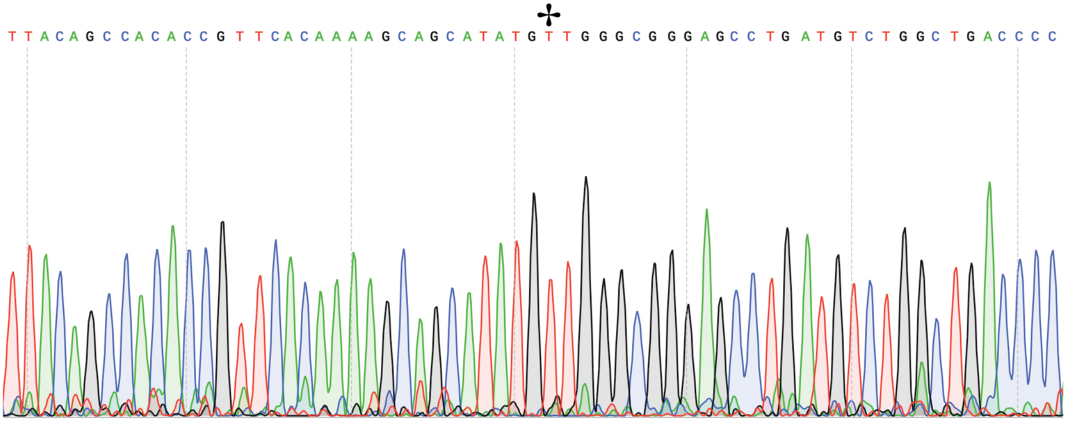


-allel 1 & 2: ACAAAAGCAGCATATGTTGGGCGGGAGCCTGATGTCTGGCTGACCC (Insert 1)

Figure S4

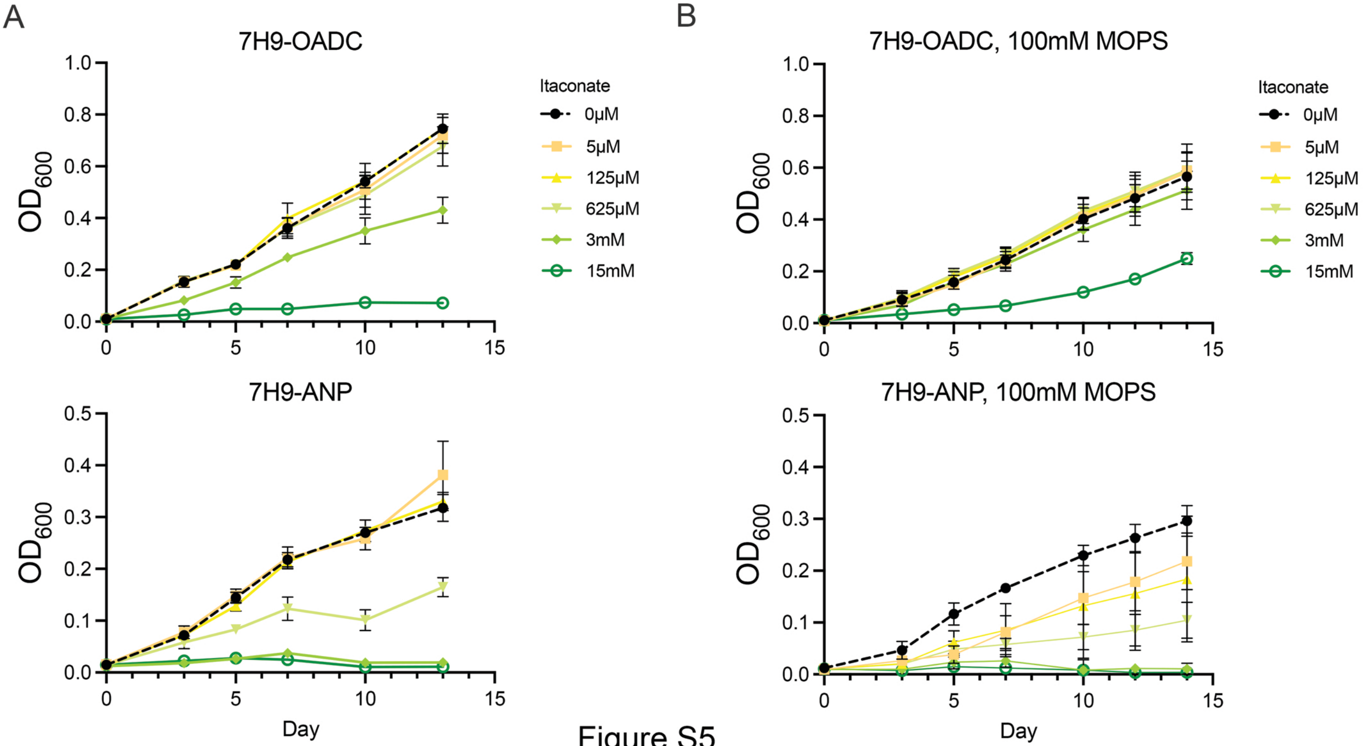
A
7H9-OADC
OD600
Itaconate
0μM
5μM
125μM
625μM
3mM
15mM
7H9-ANP
Day
B
7H9-OADC, 100mM MOPS
7H9-ANP, 100mM MOPS


Figure S5

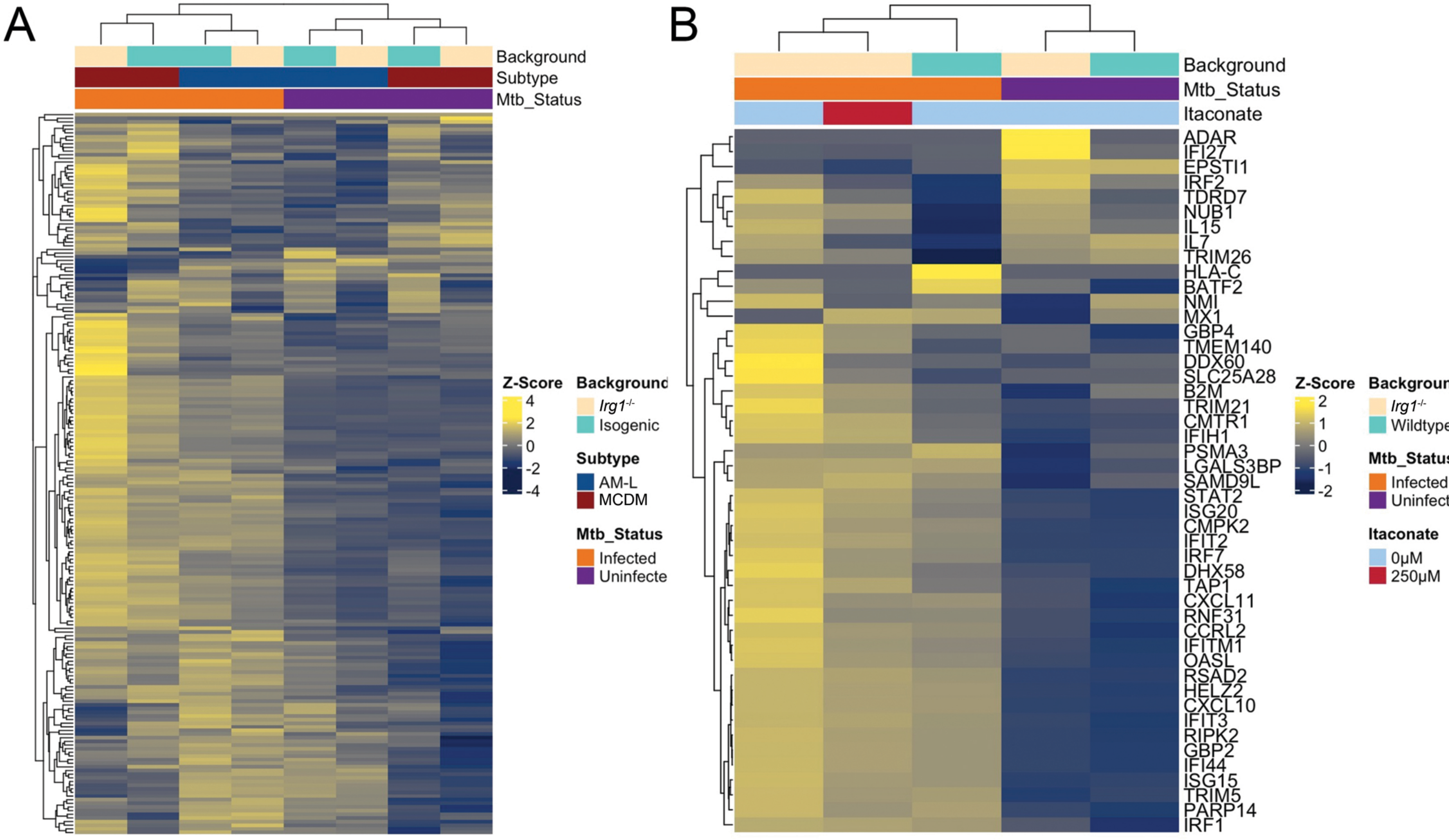

A
Background
Subtype
Mtb_Status
Z-Score
4
2
0
-2
-4
Background
Irg1-/-
Isogenic
Subtype
AM-L
MCDM
Mtb_Status
Infected
Uninfecte
B
Background
Mtb_Status
Itaconate
ADAR
IFI27
EPSTI1
IRF2
TDRD7
NUB1
IL15
IL7
TRIM26
HLA-C
BATF2
NMI
MX1
GBP4
TMEM140
DDX60
SLC25A28
B2M
TRIM21
CMTR1
IFIH1
PSMA3
LGALS3BP
SAMD9L
STAT2
ISG20
CMPK2
IFIT2
IRF7
DHX58
TAP1
CXCL11
RNF31
CCRL2
IFITM1
OASL
RSAD2
HELZ2
CXCL10
IFIT3
RIPK2
GBP2
IFI44
ISG15
TRIM5
PARP14
IRF1
Z-Score
2
1
0
-1
-2
Backgroun
Irg1-/-
Wildtype
Mtb_Status
Infected
Uninfect
Itaconate
0μM
250μM


Figure S6
